# Structural hormesis in protein aggregation: A minimal mechanistic model

**DOI:** 10.1101/2025.10.07.681066

**Authors:** Abhishek Mallela, Santiago Schnell

## Abstract

Protein aggregation underlies the pathogenesis of many neurodegenerative diseases, and inhibitors are often assumed to elicit monotonic dose–responses. We ask whether simple aggregation pathways can *intrinsically* generate hormesis—a biphasic profile with low–dose stimulation and high–dose inhibition. We formulate a minimal mechanistic model in which a single inhibitor interacts sequentially with pathway intermediates. Analysis and simulation show a robust non–monotonic response: low inhibitor doses increase aggregate formation, whereas high doses suppress it. We prove that this profile is structural—arising from chemical network topology rather than tuned kinetic parameters. The mechanism rationalizes pro–aggregating effects at low doses and underscores the need for full–range dose–response evaluation in inhibitor screening.

## 1. Introduction

Protein aggregation—the self–assembly of soluble monomers into larger, often insoluble assemblies—is central to many areas of biology and medicine. It is implicated in numerous debilitating human disorders, including Alzheimer’s and Parkinson’s diseases, and also poses challenges for the manufacturing and storage of protein therapeutics [1, 2, 3]. A persistent objective in biomedical research is therefore to regulate aggregation kinetics, often with small molecules or biological modulators that inhibit aggregation [4, 5]. In practice, efficacy is typically assessed under the implicit assumption of a monotonic dose–response; yet biological systems frequently violate this assumption [6, 7, 8].

A salient form of non–monotonicity is *hormesis*: stimulation at low doses and inhibition at high doses. Hormetic responses are widely documented across toxicology, pharmacology, ecology, and pest management [6, 8, 9], and have been framed as a “paradox of the dose” with implications for therapy and risk assessment [10]. In systems biology, such biphasic behaviour can arise from *structural* features of chemical network topology (e.g., sequential engagement of intermediates, competition for binding sites, or pathway branch points), independently of allostery, substrate inhibition, product activation, or the presence of explicit feedback and feedforward loops [11, 12].

There is growing evidence that hormesis can arise specifically in protein aggregation systems. Mild stressors such as transient heat shock or dietary restriction can reduce aggregation in models of polyglutamine diseases and *α*–synucleinopathies, whereas higher or chronic exposures are detrimental, often via HSP or Nrf2–mediated proteostasis pathways [13, 14]. Conversely, presumed inhibitors may paradoxically stimulate aspects of aggregation at low concentrations: for example, trehalose can expedite the onset of *α*–synuclein aggregation while reducing overall fibril load [15], and the small heat–shock protein *α*–crystallin can accelerate insulin aggregation under particular solution conditions [16]. In the language of our motif, such modulators act as an inhibitor engaging different intermediates with distinct affinities or activities, thereby biasing fluxes through early versus late steps in a concentration–dependent manner.

Mechanistic mathematical modeling is well–suited to disentangle these phenomena: models formulated using ordinary differential equations (ODEs) isolate network components and expose the structural features responsible for paradoxical responses [12, 17, 18]. In a related context, Rashkov et al. [17] showed that inhibition within a signaling cascade can yield hormesis in the pathway output under certain parameter regimes, underscoring the role of network competition and feedback [11]. Protein aggregation kinetics emerge from a hierarchy of coupled microscopic steps—including primary nucleation, elongation, secondary nucleation, and fragmentation—for which mean–field reductions yield tractable ODE models that successfully fit and rationalize macroscopic observables [19, 20, 21, 22, 23, 24]. These frameworks have been used extensively to interpret experiments and extract rate laws across proteins and conditions [3, 23].

In this work, we introduce a minimal phenomenological reaction mechanism of protein aggregation modulated by an inhibitor as a coarse–grained motif, abstracting a richer cascade of intermediates that represent successive pre–fibrillar species along the nucleation/oligomerisation pathway, while the mature aggregate pool collects products of elongation and secondary pathways. We make explicit the assumptions under which this coarse–graining is appropriate— early–time kinetics before substantial monomer depletion, quasi–steady behaviour of rapidly equilibrating steps, and effective rate constants that lump secondary processes. Such coarsegrained representations have proven effective for characterizing aggregation kinetics across diverse protein systems, including amyloid-*β, α*-synuclein, prions, and insulin, where phenomenological two-step models capture essential features of the aggregation time course despite abstracting over size-distribution dynamics [25, 26]. We develop a minimal mechanistic model in which a *single inhibitor* acts sequentially on pathway intermediates. Using mathematical analysis and computational simulation, we show that the model predicts a robust biphasic response of aggregate abundance over a wide range of kinetic parameters. We further *prove* that this response is structural: it follows from the topology of the reaction network rather than from finely tuned rate constants, where low inhibitor concentrations favor a productive intermediate to increase aggregate production, whereas high concentrations divert flux into an off–pathway sink and reduce aggregation. We conclude by discussing implications for inhibitor screening—including the necessity of full–range dose– response characterization—and for interpreting paradoxical low–dose stimulation in systems with aggregation.

## 2. A minimal reaction mechanism of aggregation with inhibition

We present a minimal reaction network based on well-established phenomenological principles of protein aggregation and small-molecule modulation [25, 26]. As shown in Figure 1, the reaction scheme begins with the formation of an initial complex, *C*_1_, from soluble monomeric proteins, *M* . This step represents primary nucleation, a fundamental and rate-limiting process in many aggregation pathways [2, 3]. The on-pathway progression from *C*_1_ to the final aggregate, *A*, through an intermediate state, *C*_2_, reflects the multi-step nature of aggregation, which often involves the formation of structured oligomers and protofibrils prior to the final fibrillar state.

**Figure 1.**
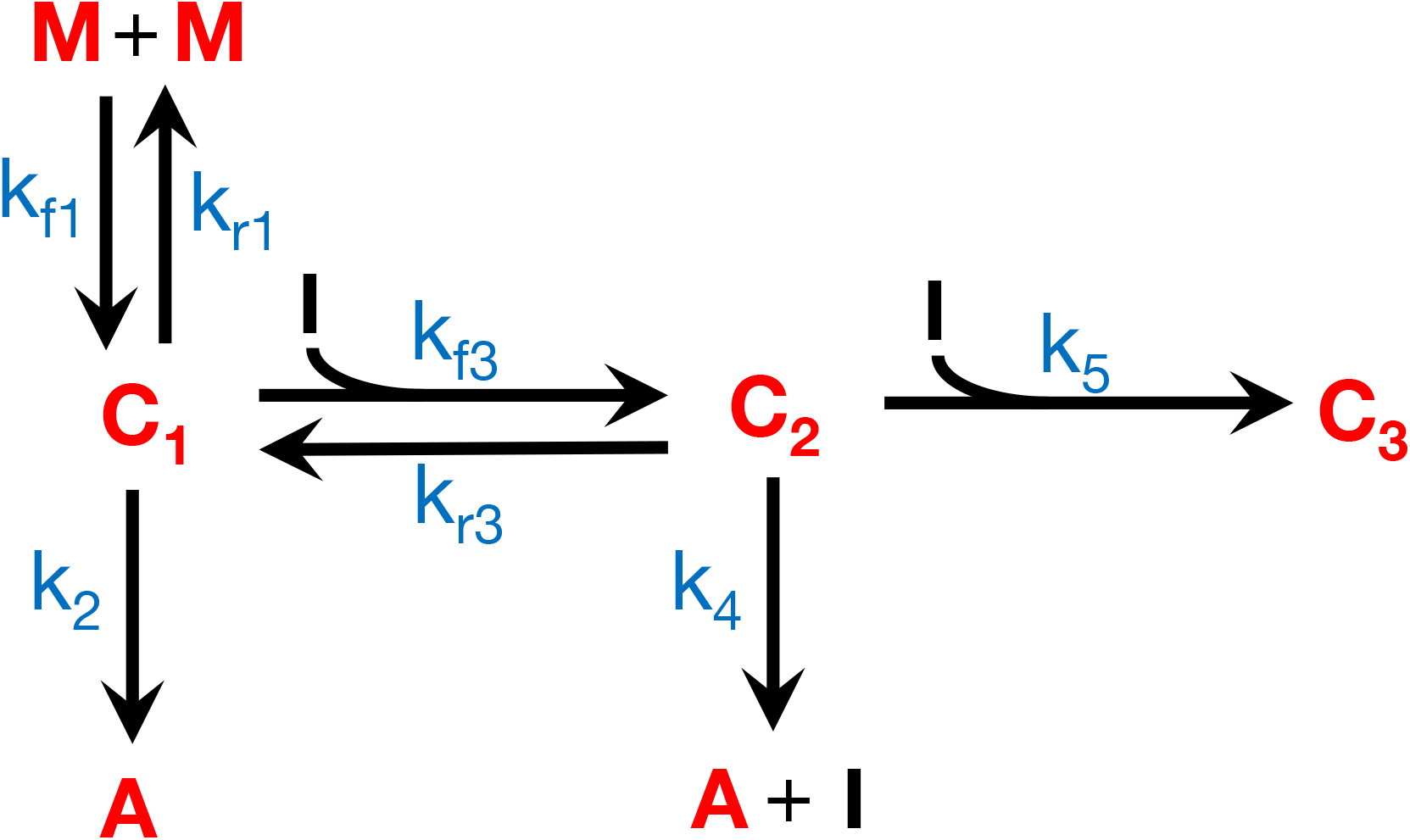
Schematic of the protein aggregation mechanism. Monomer *M* forms the dimeric intermediate *C*_1_, which converts reversibly to *C*_2_ and then to the final aggregate *A*. The inhibitor *I* interacts sequentially with intermediates, and an off–pathway sink *C*_3_ is formed at high *I* via *k*_5_. Rate constants are indicated on the arrows.

The central feature of our model is the dual role of the inhibitor (modulator), *I*. The modulator’s ability to bind the on-pathway intermediates, *C*_1_ and *C*_2_, is mechanistically plausible. At low concentrations, the modulator can promote a productive conformation or nucleation reaction step, consistent with observations of certain small molecules or chaperones that can “prime” a protein for assembly. At higher concentrations, the modulator’s interaction with the intermediate *C*_2_ leads to the formation of an inactive complex, *C*_3_.

This sequestration mechanism represents a direct and potent form of inhibition, where the modulator effectively removes productive species from the aggregation pathway.

To investigate the kinetic effects of the inhibitor in this protein aggregation mechanism, we apply the law of mass action to derive the set of ordinary differential equations governing the reactions:

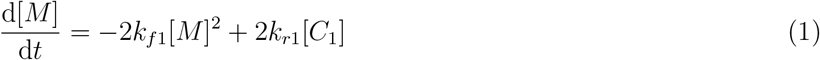

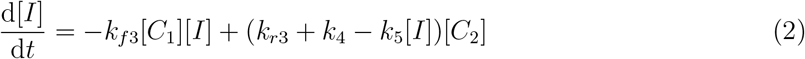

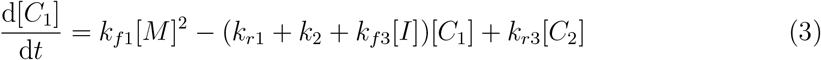

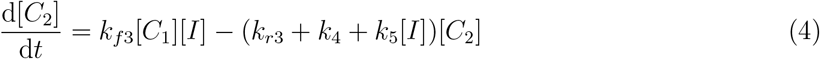

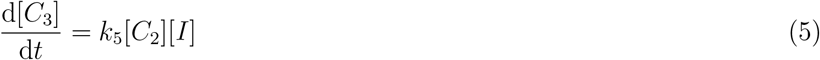

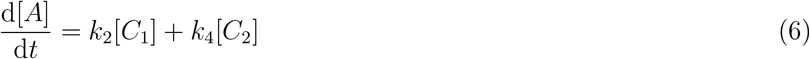

Here, [*X*] denotes the concentration of species *X*, and *k*_*i*_ are rate constants. The model’s rate constants and initial concentrations represent specific physical processes. The rates *k*_*f*1_ and *k*_*r*1_ describe the forward and reverse reactions of primary nucleation. The parameters *k*_2_ and *k*_4_ govern the rate at which intermediate species transition to the final aggregate state. The core of the modulator’s activity is captured by *k*_*f*3_, which represents its binding rate to the *C*_1_ complex, and *k*_5_, which represents its rate of sequestration into the inactive *C*_3_ sink.

### 2.1. Model simplification and non–dimensionalization

To analyze the system of ODEs, we simplified it through the techniques of timescale separation and nondimensionalization. To nondimensionalize the system, we used Kruskal’s Principle of Minimum Simplification [27]. The core idea behind this principle is to choose scaling factors that simplify the least, maintaining a maximal set of comparable terms [28].

By invoking the quasi-steady state approximation (QSSA), we set the time-derivatives of the intermediate complexes to zero: d[*C*_1_]*/*d*t* = 0 and d[*C*_2_]*/*d*t* = 0. This approximation is valid given that the intermediate species are highly reactive and their concentrations change on a much faster timescale than the primary reactants, *M* and *I* [29, 30]. *To simplify the analysis, we also applied a pseudo-first-order approximation (PFO), assuming the primary reactants’ concentrations were not significantly depleted ([M* ] ≈ *M*_*T*_, [*I*] ≈ *I*_*T*_, where *T* denotes the total concentration) [31, 32]. The approximations [*M* ] ≈ *M*_*T*_ and [*I*] ≈ *I*_*T*_ are reasonable for *in vitro* experiments where the total amount of protein and modulator is typically much larger than the concentration of reactive intermediates. The approximation [*I*] ≈ *I*_*T*_ holds true under two main conditions: either *I* is in large excess, or the binding affinity of *I* to form complexes is weak.

To demonstrate this claim, let us write down the mass conservation equation for species [*I*]. We have that the total concentration of inhibitor is equal to the sum of its concentrations in free and bound forms:

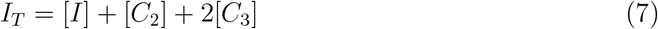

For the approximation [*I*] ≈ *I*_*T*_ to be valid, the concentration of bound *I* must be negligible compared to the total:

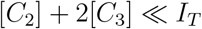

This condition is met in the following regimes:

1. **Large Excess of** *I* (*I*_*T*_ ≫ *M*_*T*_ ): The total amount of complexes that can possibly form ([*C*_1_], [*C*_2_], [*C*_3_]) is limited by the total amount of *M*, denoted *M*_*T*_ . If *I*_*T*_ ≫ *M*_*T*_, there simply is not enough *M* to create enough complexes to significantly deplete the large pool of free *I*.
2. **Weak Binding or Slow Sequestration (Large** *K*_*D*_): Even if *I*_*T*_ is not in large excess, [*I*] can remain close to *I*_*T*_ if the complexes are unstable or form slowly. If *K*_*D*_ = *k*_*r*3_*/k*_*f*3_ is large (weak binding), *C*_2_ will remain small. If [*C*_2_] is small, the rate of formation of *C*_3_ (given by *k*_5_[*C*_2_][*I*]) will also be low, keeping [*C*_3_] small, especially on fast timescales.

The parameter regimes explored in our computational experiments are consistent with these conditions, particularly at high inhibitor concentrations (where the large excess condition applies) and low concentrations (where the amount of bound complex is minimal).

We can introduce dimensionless variables by scaling concentrations with their total values (*M*_*T*_, *I*_*T*_ ) and scaling time with the characteristic rate of monomer dimerization:

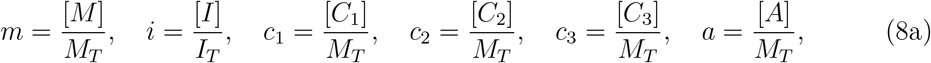

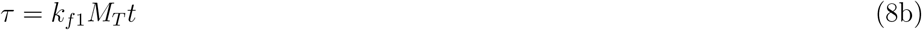

This yields a set of dimensionless parameters expressed as ratios of rate constants and initial concentrations:

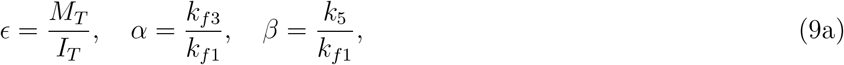

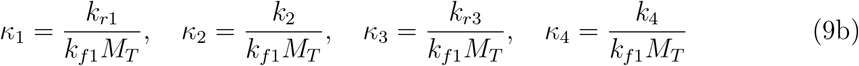

Under the QSSA, we set d*c*_1_*/*d*τ* = 0 and d*c*_2_*/*d*τ* = 0, which yields the following algebraic system for the steady-state concentrations of the intermediates:

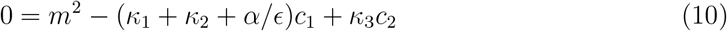

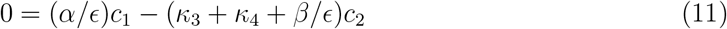

Note that the above expressions, derived under the QSSA and nondimensionalization, depend on the approximation *i* ≈ 1 (i.e., [*I*] ≈ *I*_*T*_ ).

From equation (11), we can solve for *c*_2_ in terms of *c*_1_:

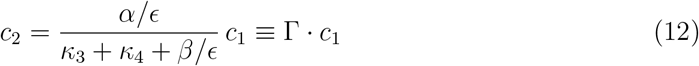

where we define

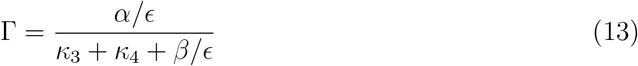

Substituting equation (12) into equation (10) gives:

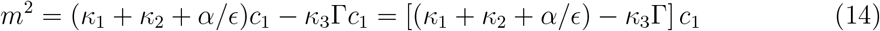

Solving for *c*_1_, we obtain:

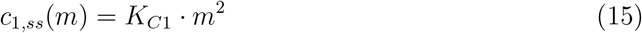

where

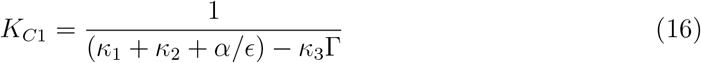

Using equations (12) and (15), we find:

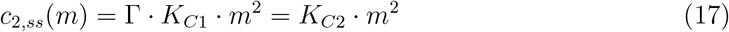

where

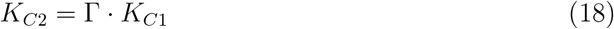

The crucial insight from the analysis above is that the steady-state levels of the intermediates are directly proportional to the square of the free monomer concentration. The composite constants *K*_*C*1_ and *K*_*C*2_ encapsulate all the kinetic parameters and concentration ratios of the system.

This process reduces the system of differential equations for the intermediates to a solvable *algebraic* system, which allows us to express their dimensionless steady-state concentrations as functions of the monomer concentration *m*, as given by equations (15) and (17).

### 2.2. Steady states and dose-response

The ODE for the monomer concentration is of the form d*m/*d*τ* = −*k*_*e*_*m*^2^, where *k*_*e*_ = 2 − 2*κ*_1_*K*_*C*1_. The solution for the dimensionless monomer concentration is:

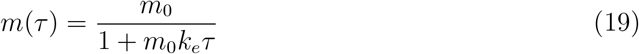

Here, *m*_0_ denotes the initial monomer concentration. Next, we substitute the solution for *m*(*τ* ) into the ODE for *a*. Let *k*_*a*_ = *κ*_2_*K*_*C*1_ + *κ*_4_*K*_*C*2_. The final solution for the dimensionless aggregate concentration is:

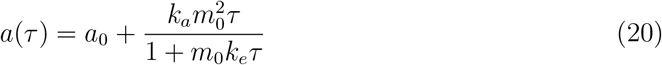

### 2.3. Proof of structural hormesis

We proved that hormesis is a structural property of the model by analyzing the quasisteady state rate of aggregation, *R* = d[*A*]*/*d*t*, as a function of the total inhibitor concentration, *I*_*T*_ . The analytical proof was performed with dimensional variables because it provides a more direct and intuitive link to the physical rate constants. Solving the algebraic system under the QSSA gives *R*(*I*_*T*_ ) as the following rational function:

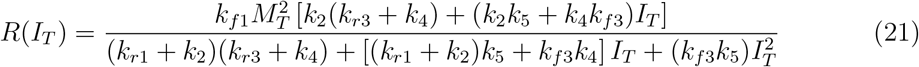

The analysis proceeded in three steps:

1. **Limit as** *I*_*T*_ → 0: In the absence of the modulator, the rate approaches a positive constant value, *R*(0) *>* 0.
2. **Limit as** *I*_*T*_ → ∞: At high modulator concentrations, the numerator is of order *I*_*T*_ while the denominator is of order 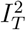. Therefore, the rate approaches zero: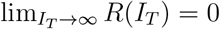.
3. **Initial Slope:** For hormesis to be manifest, the rate must initially increase. The derivative of a rational function *R*(*x*) = *N* (*x*)*/D*(*x*) at *x* = 0 is positive if *N*^*′*^(0)*D*(0) − *N* (0)*D*^*′*^(0) *>* 0. For our rate *R*(*I*_*T*_ ), this term is:

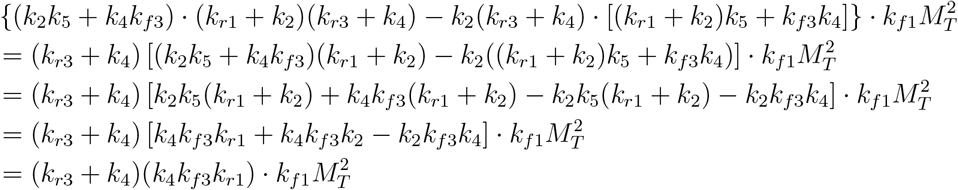

Since all rate constants (*k*_*i*_) are positive real numbers, the initial slope is strictly positive. Because the rate function starts at a positive value, has a positive initial slope, and decays to zero at infinity, it is guaranteed to exhibit a maximum for *I*_*T*_ *>* 0. This proves that hormesis is a robust, structural property of the model, independent of specific parameter values. Although this proof analyzes the quasi-steady state rate of aggregation, our computational experiments in the next section focused on the final steady-state concentration of aggregate, [*A*], which represents the most biologically relevant outcome (i.e., total pathological load).

## 3. Computational experiments

We simulated the model to determine the dose–response relationship between the total inhibitor concentration *I*_*T*_ and the total amount of aggregate *A* at steady-state. Numerical simulations showed a robust hormetic response and corroborated our analytical findings. Numerical integration of the systems was performed by the stiff solver LSODA in Python’s SciPy library. All analyses were performed at steady-state with the parameter values shown in Tables 1 and 2. The parameter values for Figure 4A were taken from Rashkov et al. [17].

**Table 1:** Parameters for Figures 2 and 3.

| Parameter | Value | Units |
| --- | --- | --- |
| $k_{f1}$ | 1.11 | $\text{nmol}^{-1} \text{L}^{-1} \text{s}^{-1}$ |
| $k_{r1}$ | 8.56 | $\text{s}^{-1}$ |
| $k_2$ | 0.10 | $\text{s}^{-1}$ |
| $k_{f3}$ | 5.39 | $\text{nmol}^{-1} \text{L}^{-1} \text{s}^{-1}$ |
| $k_{r3}$ | 4.77 | $\text{s}^{-1}$ |
| $k_4$ | 8.23 | $\text{s}^{-1}$ |
| $k_5$ | 5.37 | $\text{nmol}^{-1} \text{L}^{-1} \text{s}^{-1}$ |
| $M_T$ | $5.00 \times 10^{-3}$ | $\text{nmol L}^{-1}$ |

**Table 2:** Parameters for Figure 4B.

| Parameter | Value | Units |
| --- | --- | --- |
| Base case $k_2$ | 0.10 | $\text{s}^{-1}$ |
| High $k_2$ | 1.00 | $\text{s}^{-1}$ |
| Base case $k_4$ | 8.23 | $\text{s}^{-1}$ |
| Low $k_4$ | 1.00 | $\text{s}^{-1}$ |

### 3.1. The Inhibitor Induces a Robust Hormetic Response

As shown in Figure 2, when the total inhibitor concentration (*I*_*T*_ ) was low, the final concentration of aggregate (*A*) exceeded the basal level observed in the modulator’s absence. As *I*_*T*_ increased further, this stimulatory effect reversed, and the amount of aggregate became strongly inhibited, eventually falling far below the basal level. For the parameters used in Figure 2, the basal level of aggregation (with *I*_*T*_ = 0) was approximately 0.0011 nM. At the peak of the hormetic curve (*I*_*T*_ ≈ 0.3 nM), the aggregate concentration reached approximately 0.00175 nM. This represented a 1.6-fold increase compared to the basal level. This stimulatory effect reversed as the concentration of *I*_*T*_ increased further, leading to a profound inhibition of aggregation that fell far below the basal level.

We found that specific parameter values influenced the characteristics of the hormetic response, such as the magnitude and location of the peak. For instance, a higher sequestration rate (*k*_5_) leads to a sharper decline in aggregation, shifting the inhibitory phase to a lower modulator concentration. This suggests that while the existence of the hormetic response is a structural property of the network, its specific quantitative features are tunable by kinetic parameters. This observation provides a smooth transition to our next section, where we demonstrate that the fundamental biphasic nature of the response is independent of these specific parameter values.

### 3.2. Intermediate Dynamics Drive the Hormetic Response

To understand the mechanism underlying this biphasic response, we analyzed the steady-state concentrations of the intermediate complexes *C*_1_, *C*_2_, and *C*_3_ as a function of *I*_*T*_ (Figure 3). In the absence of the modulator, the system exists primarily as the *C*_1_ complex. At low concentrations of *I*_*T*_, *C*_1_ is efficiently converted into *C*_2_, the second productive intermediate. The rise in the concentration of *C*_2_ directly corresponds to the region of stimulated aggregation shown in Figure 2. However, as *I*_*T*_ increases, the modulator’s secondary action becomes dominant: *C*_2_ is consumed to form the inactive *C*_3_ complex. At high modulator concentrations, the system’s effective mass is almost entirely sequestered in the *C*_3_ state, effectively starving the productive pathway and causing profound inhibition of aggregation. Hormesis in this system is therefore a direct consequence of the inherent trade-off between the initial formation of the productive *C*_2_ complex and its subsequent depletion into the inactive *C*_3_ sink. Note that the apparent discontinuity at *I*_*T*_ = 2 nM is a key feature representing the underlying chemical kinetics of the system. This location marks the transition from a stimulation–dominated regime to an inhibition–dominated one, as seen in Figure 2.

**Figure 2.**
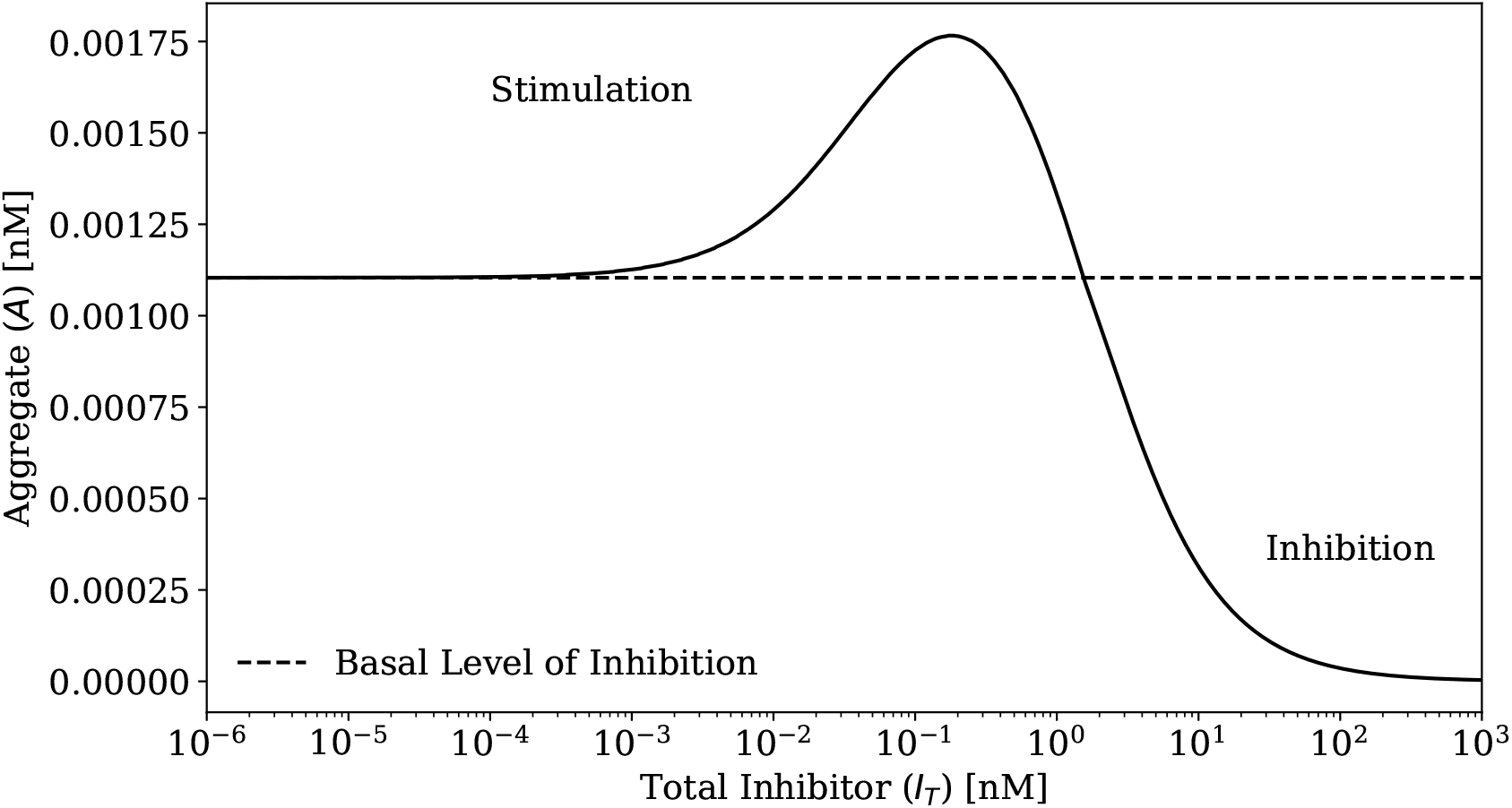
Hormetic response of the aggregate to the inhibitory stimulus. Dose–response of aggregate *A* at steady state as a function of total inhibitor *I*_*T*_ (logarithmic scale). Low doses stimulate aggregation, whereas high doses inhibit it, yielding a biphasic profile. The dashed line indicates the basal aggregate level without inhibitor. The parameter values used for the numerical simulations are available in Table 1.

**Figure 3.**
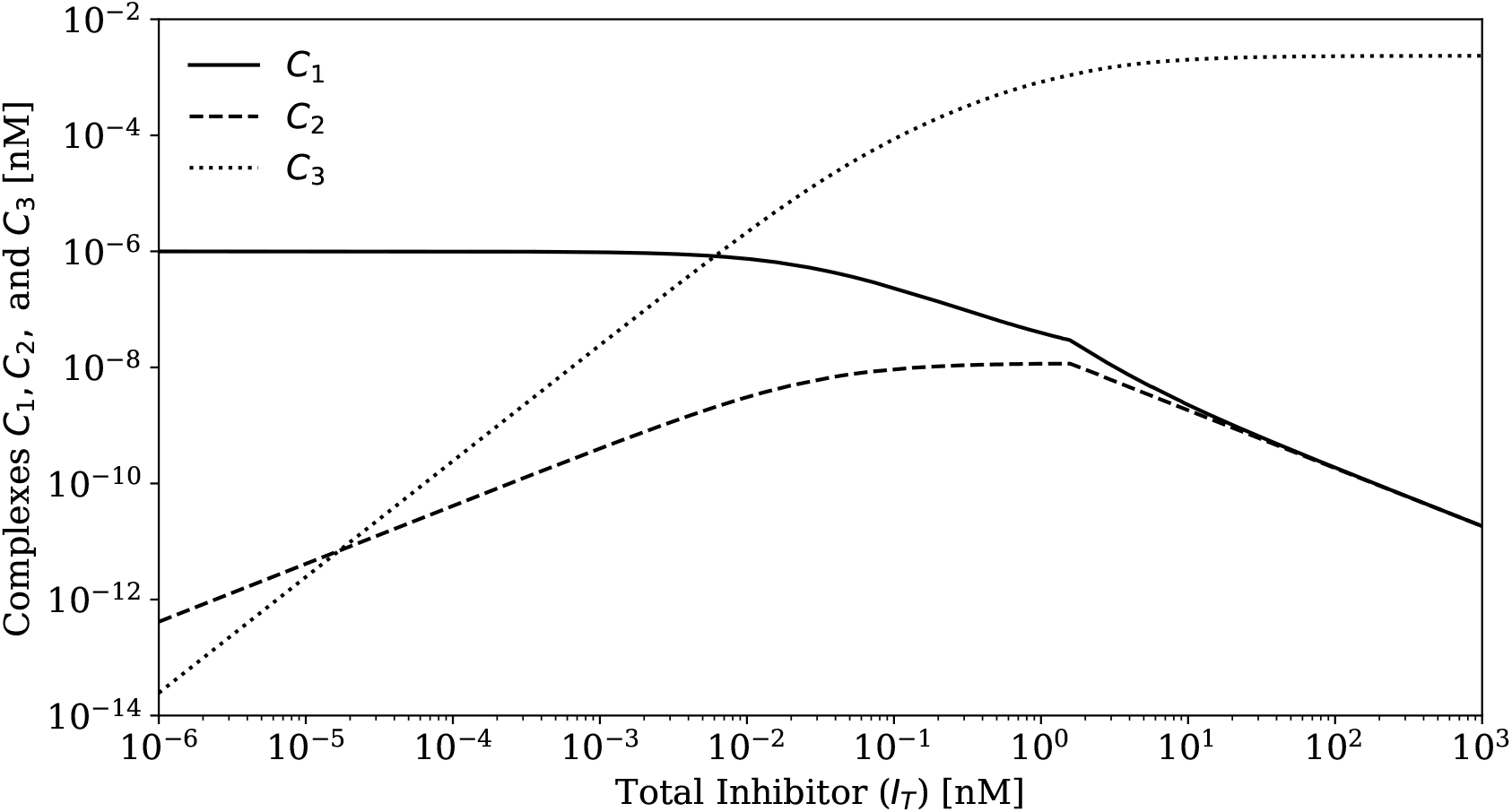
Intermediate dynamics drive the hormetic response. Intermediate complexes at steady state versus total inhibitor *I*_*T*_ (logarithmic axes). The rise of *C*_2_ underlies the stimulatory portion, while the growth of the sink *C*_3_ at high *I*_*T*_ depletes *C*_1_/*C*_2_ and produces the inhibitory portion. The parameters values used for numerical simulations are available in Table 1.

**Figure 4.**
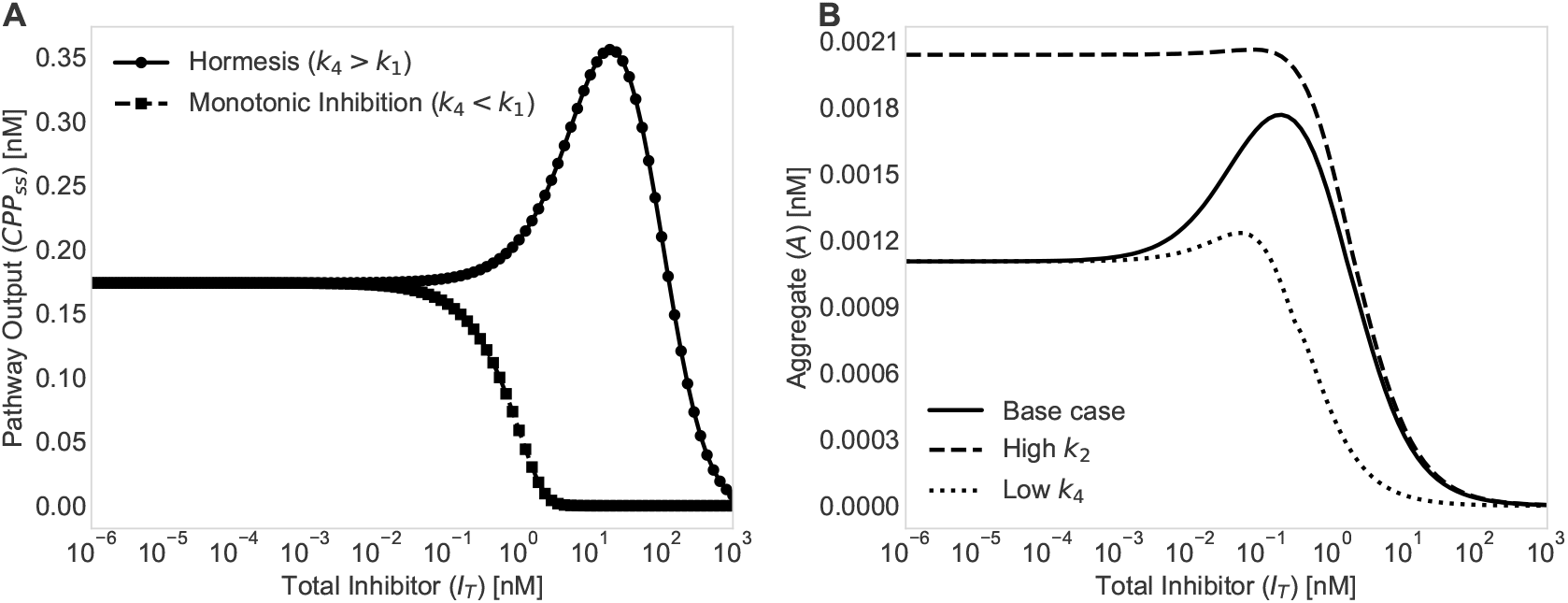
Parameter–dependent versus structural hormesis. (A) Simulation of the model by Rashkov et al. [17], where hormesis is parameter-dependent and conditional on specific kinetic parameters. Pathway output exhibits hormesis when *k*_4_ *> k*_1_ and monotonic inhibition when *k*_4_ *< k*_1_. (B) The biphasic shape of *A* versus *I*_*T*_ persists under order–of–magnitude perturbations of *k*_2_ and *k*_4_; peak location and amplitude shift but the response remains non–monotonic (simulations were carried out using the parameter sets in Tables 1 and 2). The biphasic hormetic response is conserved in all cases, confirming that it is an intrinsic structural property of the network topology.

### 3.3. Hormesis is an Intrinsic Structural Property of the Model

Finally, our analysis indicated that the observed hormetic response is a general feature of the model. In some biochemical systems, hormesis is parameter-dependent, emerging only when specific kinetic conditions are met (Figure 4A). To confirm our analytical result that this is not the case for our model, we performed simulations across a wide range of parameter sets, varying key rate constants by an order of magnitude. As shown in Figure 4B, the hormetic response was conserved in all cases. Although the magnitude and position of the peak response shifted, the fundamental biphasic shape remained. This result demonstrated that hormesis in this model is not a parameter-dependent phenomenon but rather an intrinsic and robust structural property of a network in which a modulator sequentially facilitates and then sequesters pathway intermediates. These findings present a novel mechanistic basis for hormesis in protein aggregation and caution that potential therapeutic inhibitors may have counterintuitive, pro-aggregating effects at low doses, underscoring the critical need for comprehensive dose–response evaluations in drug development and motivating targeted experimental studies to validate the proposed mechanism.

A comprehensive analysis of the parameter conditions governing the magnitude of the hormetic effect, including a phase diagram spanning three orders of magnitude in (*k*_2_, *k*_4_) space, is provided in Appendix Appendix A (see particularly Section Appendix A.3 and Figure A.5).

## 4. Discussion

In this study, we developed and analyzed a mechanistic model of protein aggregation to investigate how the process is affected by a inhibitor. Our central finding is that this minimal, plausible reaction network for protein aggregation inherently produces a robust hormetic response. We demonstrated analytically that low concentrations of a inhibitor will paradoxically accelerate the rate of aggregation, while high concentrations are inhibitory. Crucially, this biphasic behavior is not contingent on a sensitive balance of kinetic parameters but is an intrinsic property of the model’s structure, suggesting that hormesis may be a more common, yet overlooked, feature of protein aggregation kinetics than currently appreciated.

The origin of hormesis in our model can be traced to the dual role of the modulator, At low concentrations, its primary effect is to bind the initial complex *C*_1_ and catalyze its conversion to the intermediate complex *C*_2_. Since both *C*_1_ and *C*_2_ are on-pathway to forming the final aggregate *A*, and *C*_2_ can be a productive species itself, this initial step can increase the overall flux through the aggregation pathway. However, as the concentration of *I* increases, its second function becomes dominant — the binding of the productive complex *C*_2_ to form the irreversible, off-pathway complex *C*_3_. This second binding event effectively sequesters material from the aggregation pathway, leading to potent inhibition at high doses. The biphasic response is therefore a direct consequence of this trade-off, where each effect dominates in a different concentration regime.

While our work builds upon previous studies of hormesis in biochemical networks, it also presents a key distinction. For example, a similar ODE-based approach has been used to show that kinase inhibition could lead to hormesis in a signaling cascade [17]. However, in their model, the emergence of hormesis was conditional, requiring a specific inequality between two rate constants. Our model reveals a different principle: that certain network topologies are inherently poised to generate a hormetic response. The sequential activation-inactivation motif in our model guarantees a biphasic response, suggesting that the structure of the network itself can be the primary determinant of such complex dynamics. This approach highlights the power of minimal, mechanistic models to reveal fundamental principles governing complex biological phenomena. By stripping away extraneous details, we can isolate the core network topology responsible for a specific behavior. This methodology, rooted in systems biology, is broadly applicable and can be used to investigate other paradoxical behaviors in diverse biological systems, from metabolic pathways and gene regulatory networks to enzyme kinetics. Furthermore, while broader studies have characterized the general nature of hormesis across many biological systems [7], our study provides a specific, bottom-up molecular mechanism that could plausibly underlie this phenomenon in the critical context of protein aggregation. Related network-level insights have also been observed in enzymatic systems showing biphasic dose–responses due to feedback control or substrate inhibition [7].

The implications of these findings for medicine and biotechnology are significant. The most profound consequence relates to the development of therapeutic inhibitors for neurodegenerative diseases characterized by protein aggregation. Our model predicts that a compound identified as a potent inhibitor in high-throughput screens (which are often performed at high concentrations) could paradoxically exacerbate the disease pathology at lower, more physiologically relevant doses. This highlights a potential pitfall in drug discovery and underscores the critical necessity of characterizing dose–response relationships over a wide concentration range during preclinical development. Similar concerns have been raised in the context of drug-induced paradoxical effects observed in cancer therapies and neurological treatments [33]. Biologically, the modulator *I* could represent various factors, including small molecules, metal ions, or even molecular chaperones that have complex, concentration-dependent effects on protein conformation and assembly.

Our model is, by design, a simplified representation of a complex biochemical process. We have focused on a minimal network to isolate the core topology capable of producing structural hormesis. Key limitations of this coarse-grained approach include the representation of the aggregate pool *A* as a single species, which does not account for a distribution of aggregate sizes or for growth via monomer addition to existing aggregates [34]. Future extensions could incorporate such features, for instance by including an elongation step (e.g., *A* + *M* → *A*). Additionally, limitations include the absence of competing cellular pathways, such as protein degradation by the ubiquitin-proteasome system. The model also assumes a well-mixed system and does not account for the complex spatiotemporal dynamics that can occur within a cell. Spatial heterogeneity and compartmentalization have been shown to influence aggregation kinetics *in vivo* [3, 35]. Future work should focus on extending the model to incorporate these additional layers of biological reality. The critical next step, however, is experimental validation. We hope our theoretical predictions will motivate experimental studies to systematically measure aggregation rates for proteins like amyloid-beta or alpha-synuclein across a broad spectrum of concentrations for various known modulators, to determine if the intrinsic hormetic response we describe here can be observed *in vitro* or *in vivo*.

To specifically test our model’s predictions, we propose a dose–response study using the small molecule resveratrol to modulate amyloid-beta (A*β*) aggregation. Resveratrol’s effects on A*β* have been shown to be complex and concentration-dependent, including inhibiting oligomeric cytotoxicity without preventing oligomer formation [36] and even mediating the cleavage of the peptide into smaller fragments [37]. A straightforward *in vitro* experiment could be performed by incubating the A*β* protein with a wide range of resveratrol concentrations and then using a common fluorescent assay, such as Thioflavin T (ThT), to measure the amount of aggregate formed [38]. The direct binding of resveratrol to both monomeric and fibrillar A*β* has been previously demonstrated [39]. Our model predicts that the resulting dose–response curve would be bell–shaped, demonstrating a hormetic effect where low concentrations of resveratrol increase aggregation, while higher concentrations inhibit it. This outcome would provide direct experimental evidence for the proposed activation–inactivation mechanism.

## Data accessibility

All Python code and simulation data supporting this publication are publicly available at https://doi.org/10.5281/zenodo.18370730.

## CRediT Authors’ contributions

**Abhishek Mallela:** Data curation, Formal analysis, Investigation, Methodology, Visualization, Writing – original draft, review & editing. **Santiago Schnell:** Conceptualization,

Funding Acquisition, Investigation, Methodology, Project administration, Resources, Supervision, Writing – review & editing.

## Competing interests

The authors declare no competing interests.

## Funding

This work was partially funded by Dartmouth.

## Appendix A. Parameter-dependent and independent hormesis

### Appendix A.1. Parameter-dependent hormesis in the model of Rashkov et al

Rashkov et al. [17] analyzed a dual phosphorylation-dephosphorylation cycle in which a kinase inhibitor can produce hormetic responses in pathway output. In their model, hormesis is a *parameter-dependent* phenomenon: the biphasic response emerges only when the dephosphorylation rate *k*_4_ exceeds the phosphorylation rate *k*_1_ (i.e., *k*_4_ *> k*_1_). When this inequality is reversed (*k*_4_ *< k*_1_), the system exhibits monotonic inhibition.

The mathematical basis for this condition can be understood as follows. In their model, the inhibitor acts on the kinase, reducing phosphorylation flux. When the rate of dephosphorylation is fast relative to the rate of phosphorylation (*k*_4_ *> k*_1_), low inhibitor concentrations create a kinetic regime in which the substrate accumulates in an intermediate phosphorylated state that contributes to pathway output, producing transient stimulation. When phosphorylation dominates (*k*_4_ *< k*_1_), inhibition of the kinase immediately reduces flux through the pathway at all concentrations.

### Appendix A.2. Structural versus practical hormesis in the aggregation model

In contrast to the model of Rashkov et al. [17], we proved analytically (Section 2.3) that the initial slope of the aggregation rate *R*(*I*_*T*_ ) with respect to inhibitor concentration is *always positive* for any choice of positive rate constants:

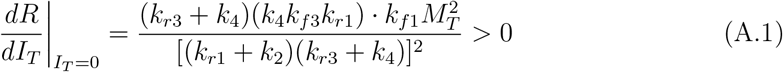

This guarantees that hormesis is a *structural* property of the network topology: the rate function must have a maximum at some *I*_*T*_ *>* 0.

However, the *magnitude* of the hormetic effect—quantified by the ratio of peak aggregate concentration to baseline aggregate concentration—depends on the relative values of the rate constants. When the direct aggregation pathway (*C*_1_ → *A* via *k*_2_) dominates the inhibitor-mediated pathway (*C*_2_ → *A* via *k*_4_), the hormetic peak becomes vanishingly small and may be practically undetectable.

### Appendix A.3. Phase diagram analysis of parameter space

To characterize the parameter regimes where hormesis is practically significant, we computed the hormesis ratio (peak aggregate concentration divided by baseline aggregate concentration) across a two-dimensional grid of *k*_2_ and *k*_4_ values spanning three orders of magnitude (10^−4^ to 10^−1^ s^−1^). Figure A.5 presents the resulting phase diagram.

The key findings are as follows:

1. **Hormetic region**: When *k*_2_ *< k*_2,*c*_, where *k*_2,*c*_ ≈ 0.045 s^−1^, the hormesis ratio substantially exceeds unity, indicating pronounced biphasic behavior. Notably, this boundary is largely independent of *k*_4_, as evidenced by the nearly vertical transition in the phase diagram.
2. **Monotonic regime**: When *k*_2_ *> k*_2,*c*_, the direct pathway dominates, and the hormesis ratio approaches unity. In this regime, the mathematically guaranteed positive initial slope produces a hormetic peak small enough as to be experimentally undetectable.

**Figure A.5:**
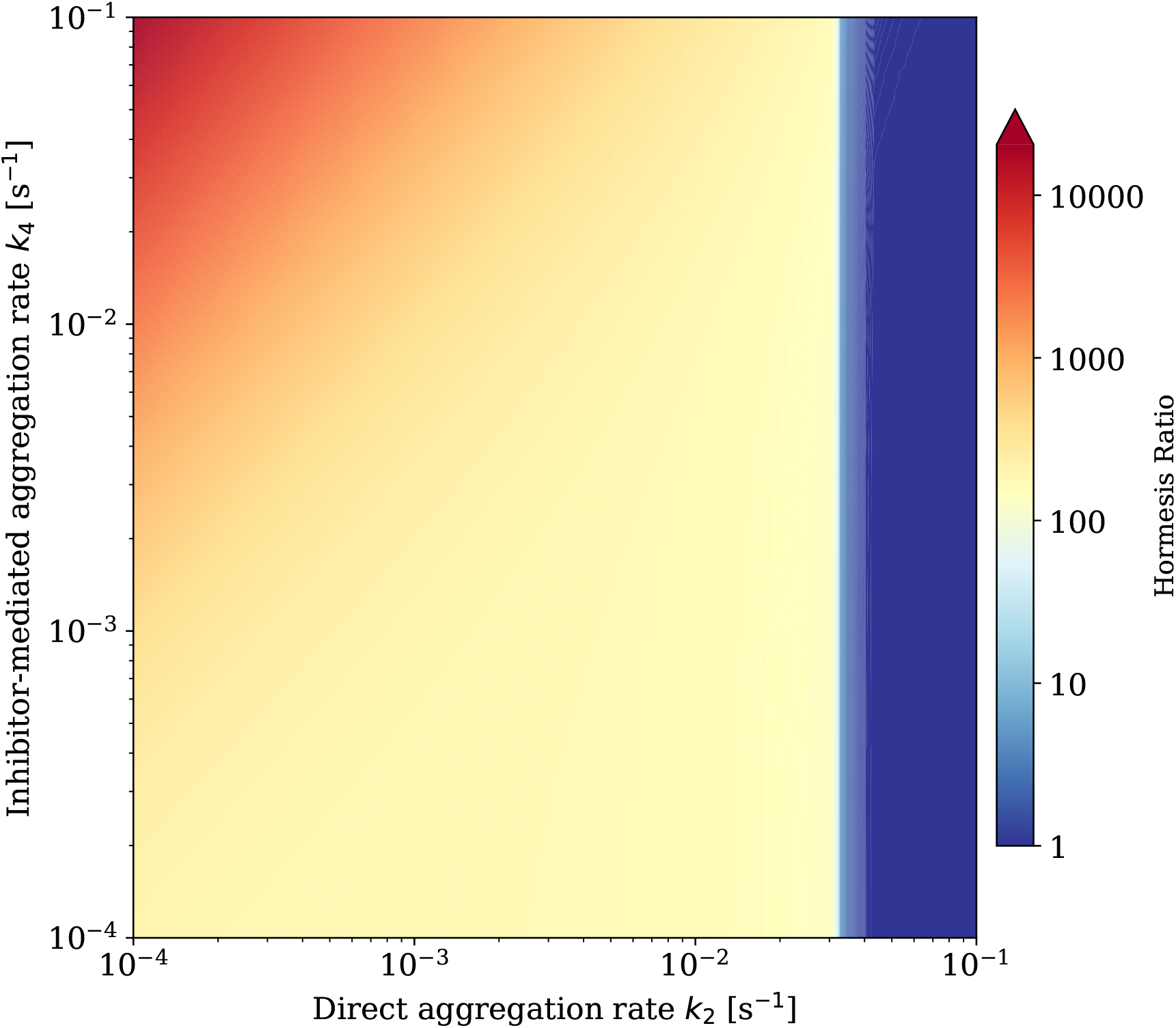
Phase diagram of hormesis. Heat map of the hormesis ratio—the maximum steady-state aggregate concentration divided by the baseline (inhibitor-free) concentration—across a 100 *×* 100 grid in the (*k*_2_, *k*_4_) parameter space. The direct aggregation rate *k*_2_ (C_1_ → A) varies along the horizontal axis; the inhibitor-mediated rate *k*_4_ (C_2_ → A) varies along the vertical axis. Both rates span from 10^−4^ to 10^−1^ s^−1^ on logarithmic scales. There is a marked transition from pronounced hormesis (warm colors, ratio *>* 1) to effectively monotonic inhibition (cool colors, ratio ≈ 1). **Methods:** For each (*k*_2_, *k*_4_) pair, the ODE system (Equations 1–6) was integrated using the stiff solver LSODA with relative and absolute tolerances of 10^−12^ for 75 logarithmically–spaced inhibitor concentrations (*I*_*T*_ = 10^−6^ to 10^3^ nM). Quasi-steady state was defined as convergence of all state variables to within a tolerance of 10^−6^ over successive 10-second integration intervals. This criterion captures the kinetically relevant quasi-steady state where hormesis manifests, rather than the true thermodynamic equilibrium where hormesis disappears. Computations were parallelized using the joblib library in Python.

### Appendix A.4. Mechanistic interpretation

The condition *k*_2_ *< k*_2,*c*_ for practical hormesis has a clear mechanistic interpretation. For the inhibitor to produce a stimulatory effect at low concentrations, the direct aggregation pathway (*C*_1_ → *A*) must be sufficiently slow that the inhibitor-mediated pathway through *C*_2_ can make a substantial contribution to aggregate formation. When the direct pathway is fast (*k*_2_ *> k*_2,*c*_), aggregate forms primarily via *C*_1_ → *A* before the inhibitor can redirect flux through *C*_2_.

The near-independence of this threshold from *k*_4_ indicates that the critical factor is not how fast *C*_2_ produces aggregate, but rather whether *C*_2_ has sufficient opportunity to form at all. When *k*_2_ exceeds the critical value, *C*_1_ is depleted so rapidly via the direct pathway that inhibitor binding cannot effectively compete, regardless of how productive the *C*_2_ → *A* step might be.

### Appendix A.5. Comparison with the model of Rashkov et al

The key distinction between the two models is summarized in Table A.3:

**Table A.3:** Comparison of parameter conditions for hormesis. With the parameter range examined in our aggregation model, the critical value *k*_2,*c*_ ≈ 0.045 s^−1^ is largely independent of *k*_4_.

| Property | Model of Rashkov et al. | Aggregation model |
| --- | --- | --- |
| Hormesis type | Parameter-dependent | Structural |
| Condition for hormesis | $k_4/k_1 > 1$ (strict) | $k_2 < k_{2,c}$ (practical) |
| Behavior when violated | Monotonic inhibition | Negligible hormesis |
| Mathematical guarantee | None | Positive initial slope |

In the Rashkov model, hormesis is a binary phenomenon—it either occurs or does not, depending on whether the parameter inequality is satisfied. In our aggregation model, hormesis is mathematically guaranteed but its practical significance depends primarily on whether *k*_2_ falls below the critical threshold *k*_2,*c*_ ≈ 0.045 s^−1^.

## Notes

https://doi.org/10.5281/zenodo.18370730

### Competing Interest Statement

The authors have declared no competing interest.

### Summary of Updates

List of changes: 1)Appendix A.3 (Phase diagram analysis): Revised the description of the hormetic and monotonic regimes to express the condition as k_2 < k_{2,c} where k_{2,c} ≈ 0.045 s^{-1}, noting that this boundary is largely independent of k_4. 2)Appendix A.4 (Mechanistic interpretation): Updated the mechanistic explanation to reflect the k_2 threshold interpretation, explaining why the condition is independent of k_4. 3)Table A.3: Updated the Condition for hormesis entry from k_4/k_2 ≳ 1 (practical) to k_2 < k_{2,c} (practical) and added clarification to the caption specifying that k_{2,c} ≈ 0.045 s^{-1}. 4)Final paragraph of Appendix A.5: Changed varies continuously with the ratio k_4/k_2 to depends primarily on whether k_2 falls below the critical threshold k_{2,c}.

